# Multidimensional trophic niche revealed by complementary approaches: gut content, digestive enzymes, fatty acids and stable isotopes in soil fauna

**DOI:** 10.1101/2020.05.15.098228

**Authors:** Anton M. Potapov, Melanie M. Pollierer, Sandrine Salmon, Vladimír Šustr, Ting-Wen Chen

## Abstract

1. The trophic niche of an organism is tightly related to its role in the ecosystem and to interactions with other species. Thousands of species of soil animals feed on detritus and co-exist with apparently low specialisation in food resource use. Trophic niche differentiation may explain species coexistence in such a cryptic environment. However, most of the existing studies provide only few and isolated evidence on food resources, thus simplifying the multidimensional nature of the trophic niches available in soil.
2. Focusing on one of the most diverse soil taxa – springtails (Collembola) – we aimed to reveal the additional value of information provided by four complementary methods: visual gut content-, digestive enzyme-, fatty acid- and stable isotope analyses, and to demonstrate the multidimensional nature of trophic niches.
3. From 40 studies, we compiled fifteen key trophic niche parameters for 125 species, each analysed with at least one method. Focusing on interspecific variability, we explored correlations of trophic niche parameters and described variation of these parameters in different Collembola species, taxonomic groups and life forms.
4. Correlation between trophic niche parameters of different methods was weak in 45 out of 64 pairwise comparisons, reflecting the complementarity of the multidimensional trophic niche approach. Gut content and fatty acids provided comparable information on fungivory and plant feeding in Collembola. Information provided by digestive enzymes differed from that gained by the other methods, suggesting its high additional value. Stable isotopes were mainly related to plant versus microbial feeding. Many parameters were affected by taxonomic affiliation but not life form. Furthermore, we showed evidence of bacterial feeding, which may be more common in Collembola than usually assumed.
5. Different methods reveal different feeding dimensions, together drawing a comprehensive picture of the trophic niche in taxa with diverse feeding habits. Food web studies will benefit from simultaneously applying several joint approaches, allowing to trace trophic complexity. Future studies on the multidimensional trophic niche may improve understanding of food-web functioning and help to explain species coexistence in cryptic environments such as soil.

## Introduction

Trophic interactions between organisms influence structure and stability of ecological communities, channeling of energy through food webs, and functioning of ecosystems (Barnes et al., 2018; Rooney & McCann, 2012). The trophic interactions of an organism with other coexisting species can be perceived as its ‘trophic niche’, stemming from the Hutchinsonian ‘ecological niche’ concept (Holt, 2009). Trophic niche can be defined in multidimensional space, where each axis represents an aspect, or dimension, of trophic interactions (Hutchinson, 1978; Machovsky-Capuska, Senior, Simpson, & Raubenheimer, 2016; Newsome, Martinez del Rio, Bearhop, & Phillips, 2007). Such dimensions could represent direct evidence of food objects, describe feeding behavior of individuals, and imply basal resources and trophic level of species in food webs with potential interactions with other species.

Soil food webs rely on dead plant and animal material, soil organic matter, phototrophic microorganisms, roots and root exudates, inseparably mixed with bacteria and fungi, and altogether generalized as ‘detritus’ (Moore et al., 2004). Thousands of soil animal species, known as ‘detritivores’ and ‘microbivores’, feed on this mixture and locally co-exist with apparently low specialisation, which inspired J.M. Anderson to formulize the ‘enigma of soil animal diversity’ (Anderson, 1975). Trophic niche differentiation is one mechanism explaining species coexistence in soil. Understanding of trophic niche differentiation in soil invertebrates, however, was for a long time constrained by the small size and cryptic lifestyle of these animals, and only few trophic niche dimensions are known. In natural soil habitats, four common methods are used, each with its advantages and drawbacks:

1. Visual gut content analysis provides reliable data on ingested food materials by microscopic observations of gut content and counting different types of particles (Anderson & Healey, 1972; Hagvar & Kjondal, 1981; Ponge, 2000). Fungal spores and hyphae in the gut indicate fungivory of soil animals, while coarse plant detritus, roots and shoots suggest herbivory and litter grazing, and amorphous material such as fine detritus may imply feeding on soil organic matter and faecal pellets. However, visual gut content analysis only presents a snapshot of the ingested materials, overestimates poorly digestible particles and provides limited information in case of feeding on fluids.
2. Digestive enzymes, such as cellulase, chitinase and trehalase, represent the ability of an animal to decompose corresponding types of organic compounds, and provide a way to assess which ingested materials may be digested (C. O. Nielsen, 1962; Parimuchová et al., 2018; Siepel & de Ruiter-Dijkman, 1993). Cellulose is a major component of cell walls of green plants and algae; cellulase activity suggests herbivory, algivory or litter grazing of soil animals. Chitin is a major component of fungal cell walls and chitinase activity suggests fungivory. Trehalose, by contrast, is a storage component of fungal, lichen and plant cells; trehalase activity can be used as a proxy for fungivory and herbivory. Furthermore, foraging strategies of soil animals can be inferred by a combination of the three digestive enzyme analyses : ‘grazers’, which can digest both cell-walls and cell-contents, have a higher activity of cellulase and chitinase to degrade structural polysaccharides, while ‘browsers’, which can only digest cell-contents, have a higher activity of trehalase to degrade storage polysaccharides (Siepel & de Ruiter-Dijkman, 1993). However, digestive enzymes provide information on potential, rather than real assimilation of food compounds.
3. Neutral lipid fatty acid (FA) analysis, by contrast, detects assimilated compounds that are retained in the fat body of consumers, a phenomenon called ‘dietary routing’ (Chamberlain, Bull, Black, Ineson, & Evershed, 2005a; Ruess & Chamberlain, 2010). Plants, fungi and different groups of prokaryotes synthesise specific membrane lipids and these compounds can be tracked in animal consumers over a period of time (usually few weeks for mesofauna; Haubert, Pollierer, & Scheu, 2011) and across trophic levels (Pollierer, Scheu, & Haubert, 2010). An extensive review of the fatty acid method for soil food web analysis can be found in Ruess & Chamberlain (2010). Despite being informative, FA analysis does not provide estimation of a species trophic level in the soil food web. Quantitative comparisons among contributions of different food origins are also limited (Kühn, Schweitzer, & Ruess, 2019).
4. Similar to FA analysis, stable isotope analysis provides information on assimilated food resources of soil animals integrated over time, but the method is quantitative and allows for trophic level estimation (Tiunov, 2007). Low ^13^C concentration in animal body tissue indicates utilisation of freshly fixed plant carbon (e.g. herbivory), while high ^13^C concentration suggests consumption of microbially processed organic matter (e.g. soil feeding, bacterivory or fungivory) (A. M. Potapov, Tiunov, & Scheu, 2019). The ^15^N concentration, by contrast, infers trophic levels of animals in the food web, being low in primary consumers but high in predators and mycorrhizal fungi feeders (A. M. Potapov, Tiunov, & Scheu, 2019). However, bulk natural stable isotopes provide only rough information on the trophic position and rarely allow to reconstruct exact feeding interactions in soil.

Different dietary methods provide information on different trophic niche dimensions of consumers over different time scales. A recent review revealed that only few studies have conducted quantitative comparisons among different methods, and none of them simultaneously applied multiple methods (J. M. Nielsen, Clare, Hayden, Brett, & Kratina, 2018). This motivated us to compile a trophic trait dataset across the four abovementioned methods from field studies and to analyse trophic niche differentiation among soil animals. We chose springtails (Collembola) as an example, since they are one of the most abundant and diverse soil invertebrates and traditionally considered as generalistic fungivores (Hopkin, 1997). However, detritivorous Collembola may assimilate only a small percentage of the ingested food (Jochum et al., 2017). Although most of Collembola are ‘herbo-fungivorous grazers’, having cellulase, chitinase and trehalase activity in digestive system (Berg, Stoffer, & van den Heuvel, 2004), they in fact occupy different trophic positions in the soil food web, spanning from algivores to high-level consumers, as indicated by the stable isotope ^15^N values (Chahartaghi, Langel, Scheu, & Ruess, 2005; Rusek, 1998). Different species of Collembola also differ in FA compositions, suggesting that they rely on food resources of different origins (Ferlian, Klarner, Langeneckert, & Scheu, 2015; T.-W. Chen, Sandmann, Schaefer, & Scheu, 2017). In particular, trophic niches of Collembola species likely correlate with taxonomic position and life form. The former correlation may suggest phylogenetic constraints, while the latter implies microhabitat specialisation in Collembola trophic niches (A. M. Potapov, Semenina, Korotkevich, Kuznetsova, & Tiunov, 2016). In this study we aimed to (1) quantitatively assess the complementarity provided by different dietary methods; (2) describe multidimensional trophic niches among different species, taxonomic groups and life forms of a model soil animal group (Collembola).

## Materials and methods

We compiled trophic data on Collembola from field studies that used visual gut content, digestive enzyme, FA and stable isotope analyses. Data were collected from the personal libraries of the authors and complemented with searching for published literature in the Web of Science. A complete list of studies can be found in Supplementary Materials. Most of the published studies applied only one method and only two used a combination of two methods (Haubert et al. 2009; Ferlian et al. 2015). For each study we averaged individual measurements by species and ecosystem for fifteen trophic niche parameters derived from the four methods (**Table 1**).

**Table 1.**
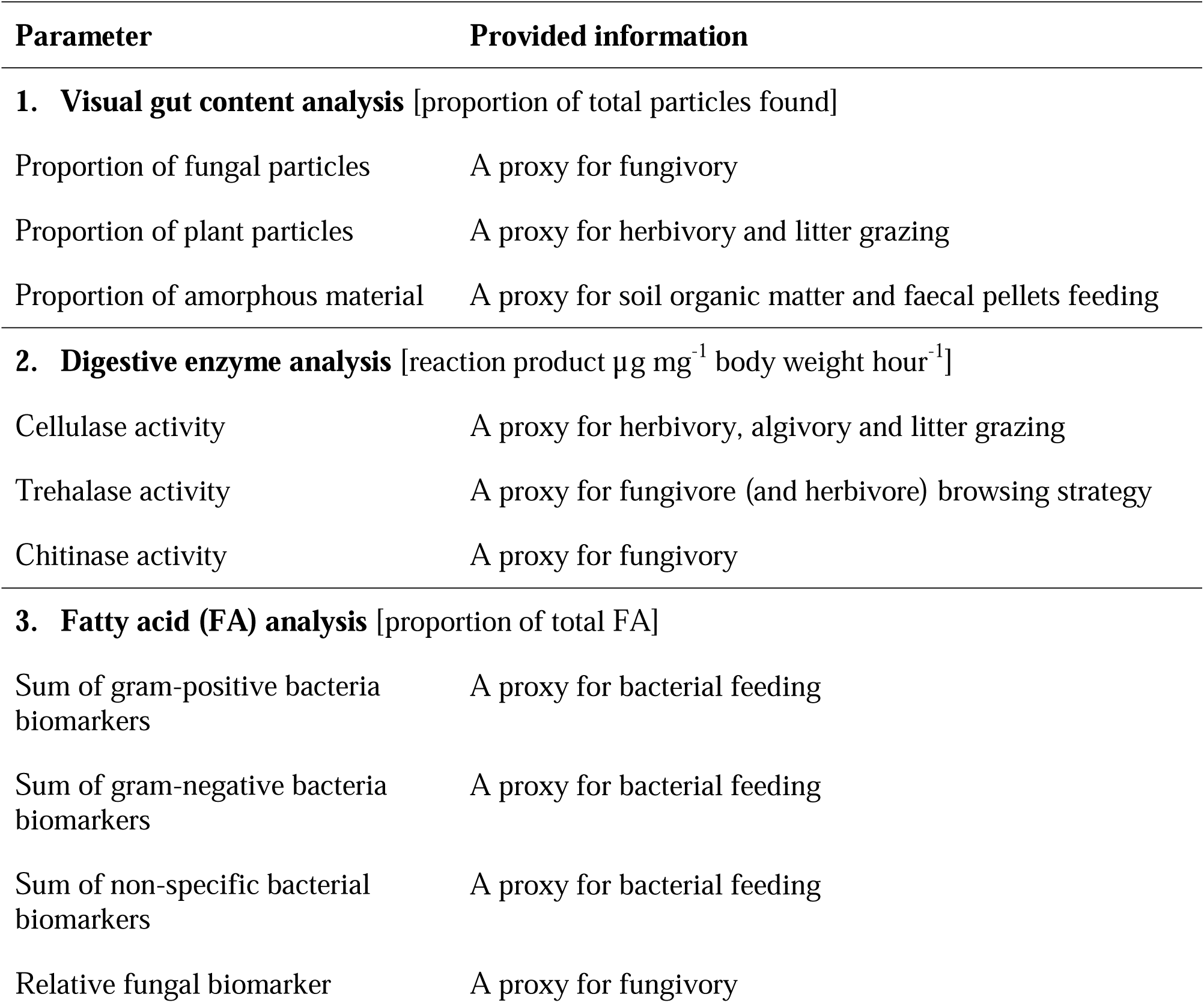
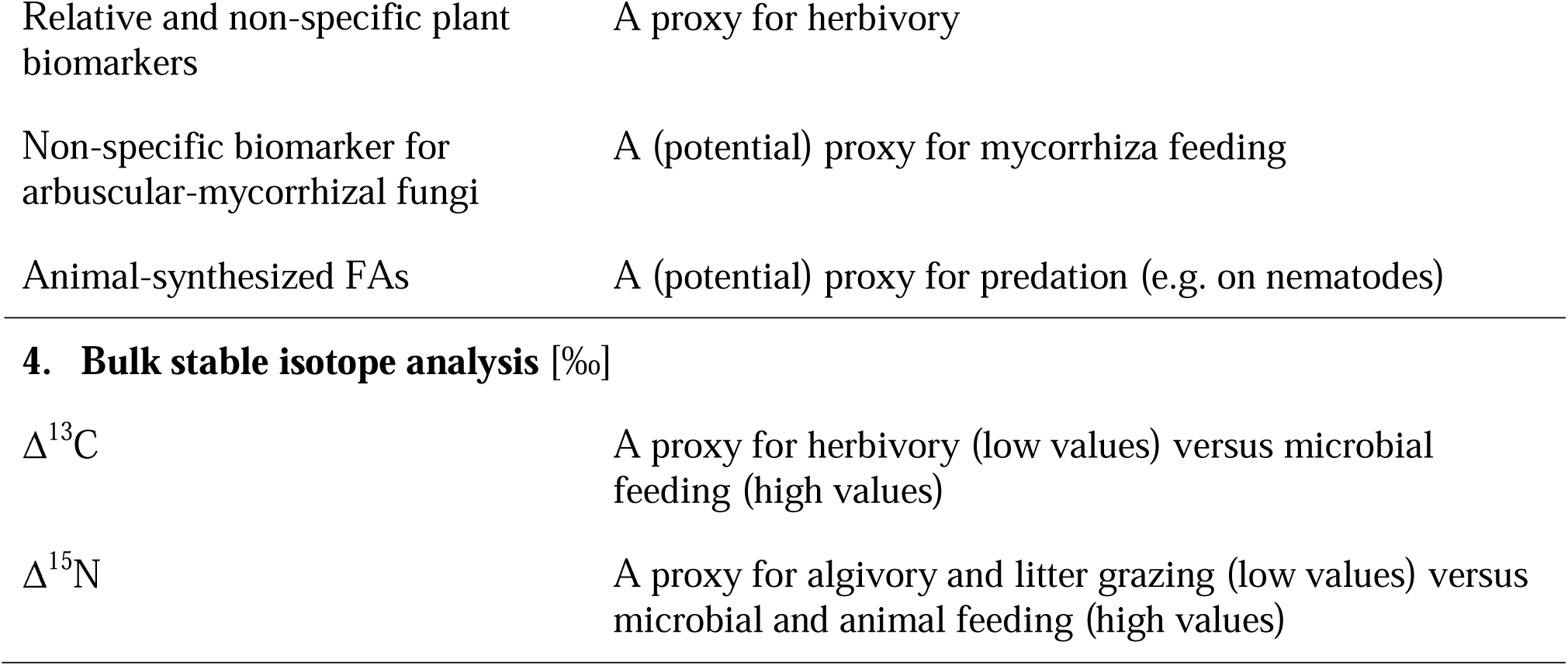
List of the trophic niche parameters used in this study. Units are given in square brackets.

| Parameter | Provided information |
| --- | --- |
| **1. Visual gut content analysis** [proportion of total particles found] |  |
| Proportion of fungal particles | A proxy for fungivory |
| Proportion of plant particles | A proxy for herbivory and litter grazing |
| Proportion of amorphous material | A proxy for soil organic matter and faecal pellets feeding |
| **2. Digestive enzyme analysis** [reaction product $\mu\text{g mg}^{-1}$ body weight $\text{hour}^{-1}$] | |
| Cellulase activity | A proxy for herbivory, algivory and litter grazing |
| Trehalase activity | A proxy for fungivore (and herbivore) browsing strategy |
| Chitinase activity | A proxy for fungivory |
| **3. Fatty acid (FA) analysis** [proportion of total FA] |  |
| Sum of gram-positive bacteria biomarkers | A proxy for bacterial feeding |
| Sum of gram-negative bacteria biomarkers | A proxy for bacterial feeding |
| Sum of non-specific bacterial biomarkers | A proxy for bacterial feeding |
| Relative fungal biomarker | A proxy for fungivory |
| Relative and non-specific plant biomarkers | A proxy for herbivory |
| Non-specific biomarker for arbuscular-mycorrhizal fungi | A (potential) proxy for mycorrhiza feeding |
| Animal-synthesized FAs | A (potential) proxy for predation (e.g. on nematodes) |
| 4. Bulk stable isotope analysis [‰] |  |
| $\Delta^{13}\text{C}$ | A proxy for herbivory (low values) versus microbial feeding (high values) |
| $\Delta^{15}\text{N}$ | A proxy for algivory and litter grazing (low values) versus microbial and animal feeding (high values) |

### Visual gut content

Primary screening of literature yielded 31 studies that reported data on the gut content of Collembola. We selected those that provided quantitative estimates from natural environments, mostly temperate forests and grasslands. We took the most commonly reported categories and defined gut content parameters as (1) particles of fungal origin, including hyphae and spores, (2) particles of plant origin, mostly coarse plant detritus (excluding pollen and algae), and (3) amorphous material of unknown nature (i.e. fine detritus such as soil organic matter). The final dataset for the three gut content parameters included 77 records on 56 species from 15 studies (**Table S**1). Raw data were expressed as proportion of certain type of particles among the total particles ingested.

### Digestive enzymes

To our knowledge, cellulase, trehalase and chitinase activities in Collembola were reported only in three studies (Berg et al., 2004; Parimuchová et al., 2018; Urbášek & Rusek, 1994). Despite using the same conceptual method, these studies used different chemical protocols and ways of glucose detection, which resulted in evident differences in absolute mean values of substrate production per unit of animal body mass. Thus, we excluded the study of (Urbášek & Rusek, 1994). The final dataset included 45 records on 27 species (**Table S2**). Raw data of digestive enzyme activity were expressed as mg of reaction products per g of animal mass per hour.

### Fatty acids

Screening of literature yielded 10 studies that reported neutral lipid FA compositions of Collembola. The dataset was complemented with unpublished data collected by Melanie M. Pollierer. Studies varied in completeness of FA profiles, but most of them reported data on 16:1ω7 and 18:1ω7 as general bacteria biomarkers, 18:2ω6,9 as relative fungal biomarker, 18:1ω9 (in addition 21:0, 22:0, 23:0, 24:0) as relative plant biomarkers and several gram-positive bacterial biomarkers (including i15:0, a15:0, i16:0, i17:0, a17:0) and gram-negative bacterial biomarkers (including 2-OH 10:0, 2-OH 12:0, 3-OH 12:0, 2-OH 14:0, 3-OH 14:0, 2-OH 16:0, cy17:0, cy19:0). If reported, we also used 16:1ω5 as the biomarker related to feeding on arbuscular-mycorrhizal fungi (Ngosong, Gabriel, & Ruess, 2012) and sum of 20:1ω9, 22:6ω3, 22:2ω6, 22:1ω9 and 24:1 as the FAs in metazoan animals (e.g. nematodes) (Chamberlain & Black, 2005; Chamberlain, Bull, Black, Ineson, & Evershed, 2005b; J. Chen, Ferris, Scow, & Graham, 2001; Tanaka et al., 1996). Raw data was compiled as proportions of the total neutral lipid FAs of the organism. Individual biomarker FAs were summed up to generate seven conventional FA parameters (**Table 1**). The final dataset included 130 records on 47 species from 13 studies (**Table S3**).

### Stable isotopes

The dataset was based on the previous compilation of Potapov et al. (2016) and complemented with data on grassland and forest communities, including recent publications. For each study, isotopic baseline (i.e. plant litter) was used to calculate litter-calibrated Δ^13^C and Δ^15^N values of Collembola (A. M. Potapov, Tiunov, & Scheu, 2019). The final dataset for the two stable isotope parameters included 378 records on 96 species from 10 studies (**Table S4**).

### Data analysis

For the fifteen trophic niche parameters derived from the four methods (**Table 1**) we in total included 125 species, each analysed with at least one method (**Table S1-4**; Dryad Digital Repository, doi:10.5061/dryad.xxxxx). All calculations were based on these datasets and conducted in R 3.5.3 (R Core Team, 2019).

Data on digestive enzyme activity were added with 0.02 μg product mg^-1^ h^-1^ (the minimum positive value observed) for all values and then log10-transformed. We normalized data of chitinase activity separately for Berg et al. (2004) and Parimuchová et al. (2018) by subtracting mean and dividing by standard deviation to account for the differences in chemical protocols. Proportional data (gut content and FA) were logit-transformed, with 0 and 1 proportions adjusted to 0.025 and 0.975, respectively, using the *logit* function in the *car* package.

Focusing on interspecific variability across trophic parameters, we conducted species-based analyses using the species names following GBIF (https://www.gbif.org/tools/species-lookup). For each trophic parameter and each species, we averaged data across ecosystems and studies. To have the same data representation across all fifteen parameters, each parameter was scaled between 0 (lowest observed value of the parameter) and 1 (highest observed value of the parameter). All the following analyses and results were based on the scaled data.

We performed three analyses: First, we tested pairwise correlations among all trophic niche parameters. Spearman rank correlation was applied using the *cor* function. The number of points for each correlation varied among parameters according to the number of shared species of the paired parameters (**Table S6**). Correlation tests were conducted for those with a minimum of six data points.

Second, we explored the association of species with their trophic parameters and visualised interspecific differentiation in multidimensional trophic niches using principal component analysis (PCA) with the *prcomp* function. We selected species according to their common trophic parameters available. Only two species had data across all fifteen parameters. Thus, we excluded parameters with fewer species, and finally chose six species that had data across seven parameters, with a focus on gut content and fatty acid parameters.

Third, we tested the effect of taxonomic affiliation (as the proxy of phylogenetic group) and life form (as the proxy of microhabitat preference) on each of the fifteen trophic niche parameters using linear models (the *lm* function). Groups with fewer than three species were excluded from the analysis. Species of Dicyrtomidae, Katiannidae, Sminthuridae, Sminthurididae, Bourletiellidae and Arrhopalitidae and species of Onychiuridae and Tullbergiidae were pooled at a higher taxonomic rank, Symphypleona and Onychiuroidea, respectively, due to low number of species in each family and trophic similarity among families (A. M. Potapov, Semenina, et al., 2016). Definition of life form followed Gisin (1943) as interpreted by A. M. Potapov, Semenina, et al., (2016) and species were categorised into atmobiotic (aboveground and surface dwellers), epedaphic (surface and upper litter dwellers), hemiedaphic (litter dwellers) and euedaphic (lower litter and soil dwellers). We further divided the best replicated family Isotomidae in epedaphic versus hemiedaphic and euedaphic species to assess the effect of life form on trophic niche parameters within this family. We reported R^2^ and p values from the model output to compare predictability of different trophic niche parameters among taxonomic groups and among life forms. For each parameter we reported median values for taxonomic groups and life forms. We then used PCA to visualise multi-dimensional trophic niches among taxonomic groups.

## Results

### Correlation between trophic niche parameters

In 45 out of total 64 performed pairwise tests (excluding within-method tests), correlation between trophic niche parameters was weak or absent (R^2^ < 0.15; **Fig. 1**; for more details see the Supplementary **Fig. S1**). Among strong positive correlations, species with high proportion of fungal and plant particles in their guts also had high proportions of fungal-synthesised (R^2^ = 0.56) and plant-synthesised FAs (R^2^ = 0.50), respectively. A high proportion of fungal particles in the gut, however, negatively correlated with retained FA synthesised by gram-negative bacteria (R^2^ = 0.46).

**Figure 1.**
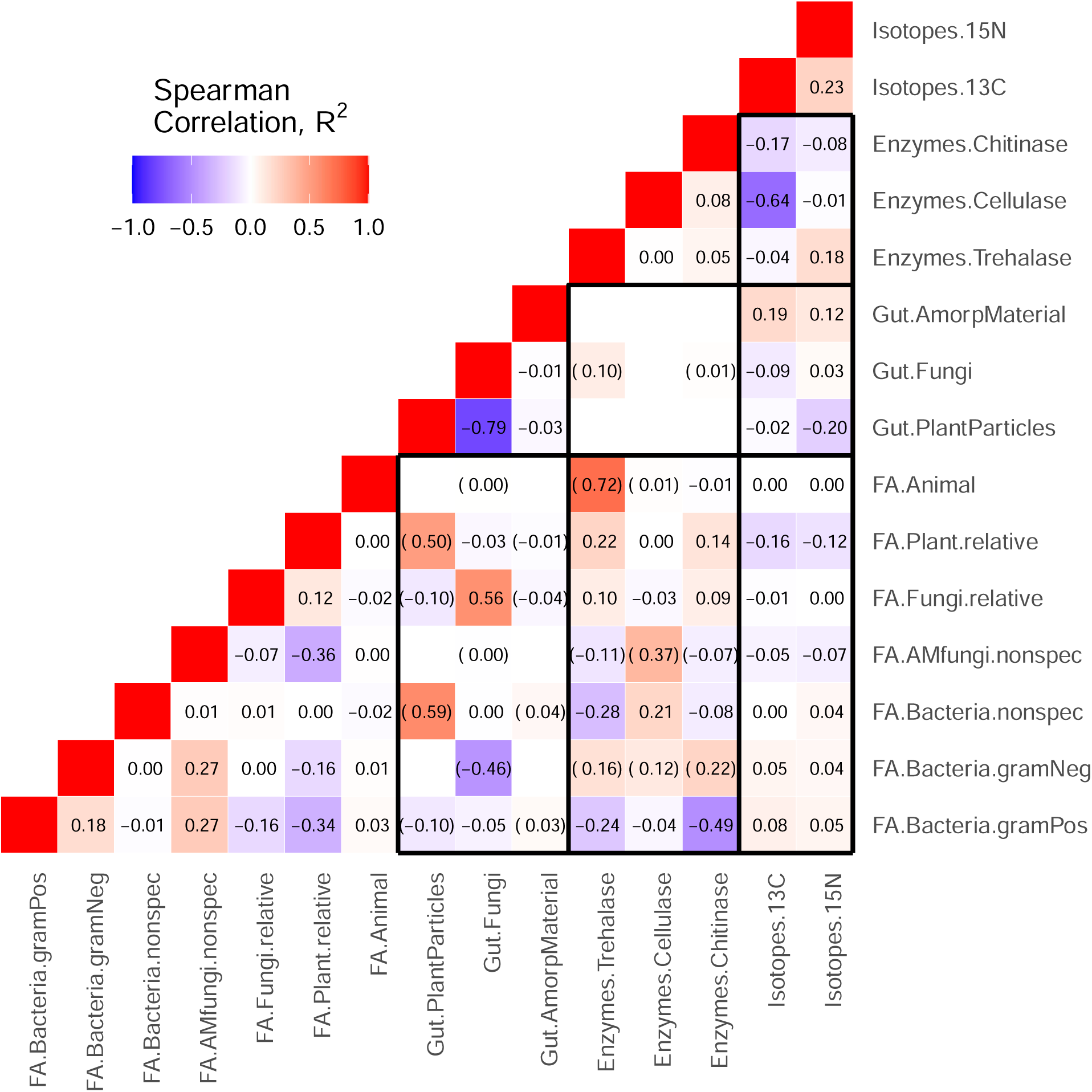
Correlation among fifteen trophic niche parameters in Collembola. Red colour represents positive correlations, blue colour represents negative correlations; to display negative correlations R^2^ was multiplied by -1. Correlations based on < 6 species were excluded, correlations based on 6–9 species are given in brackets; in other cases, n = 10–88 (**Table S5**). Correlations among parameters derived from different methods are framed within black rectangles.

Trehalase activity in guts positively correlated with the proportions of retained FAs synthesized by fungi (R^2^ = 0.10), gram-negative bacteria (R^2^ = 0.16), plants (R^2^ = 0.22) and animals (R^2^ = 0.72) in the bodies, but negatively with the proportions of FAs synthesised by gram-positive bacteria (R^2^ = 0.24) and non-specific bacteria biomarkers (R^2^ = 0.28). Cellulase activity positively correlated with the proportion of arbuscular-mycorrhizal fungi (R^2^ = 0.37) and non-specific bacteria FAs (R^2^ = 0.21). Similar to trehalase activity, chitinase activity negatively correlated with the proportion of FAs synthesised by gram-positive bacteria (R^2^ = 0.49) and positively with the proportion of FAs synthesised by gram-negative bacteria (R^2^ = 0.22).

The Δ^13^C values negatively correlated with cellulase activity (R^2^ = 0.64), chitinase activity (R^2^ = 0.17) and proportion of plant-synthesised FAs (R^2^ = 0.16). The Δ^15^N values negatively correlated with proportion of plant particles in gut (R^2^ = 0.20) and plant-synthesised FAs (R^2^ = 0.12), but positively with trehalase activity (R^2^ = 0.18).

### Multidimensional trophic niches of Collembola species

The strongest distinction in multidimensional trophic niche space was observed between *Orchesella flavescens* (surface-dwelling species of Entomobryidae) and *Protaphorura* (soil-dwelling genus of Onychiuridae; **Fig. 2**). The former species was associated with high proportions of plant-synthesised FAs and plant particles in the gut, and the latter with high proportions of fungi-synthesised FAs and fungi in the gut. *Tomocerus minor* (Tomoceridae) was associated with plant and non-specific bacteria parameters. *Lepidocyrtus lignorum* (litter-dwelling species of Entomobryidae) was related to fungi in the gut. *Parisotoma notabilis* (litter-dwelling species of Isotomidae) and *Onychiurus* (soil-dwelling genus of Onychiuridae) were associated with gram-positive bacteria FAs and amorphous material in the gut.

**Figure 2.**
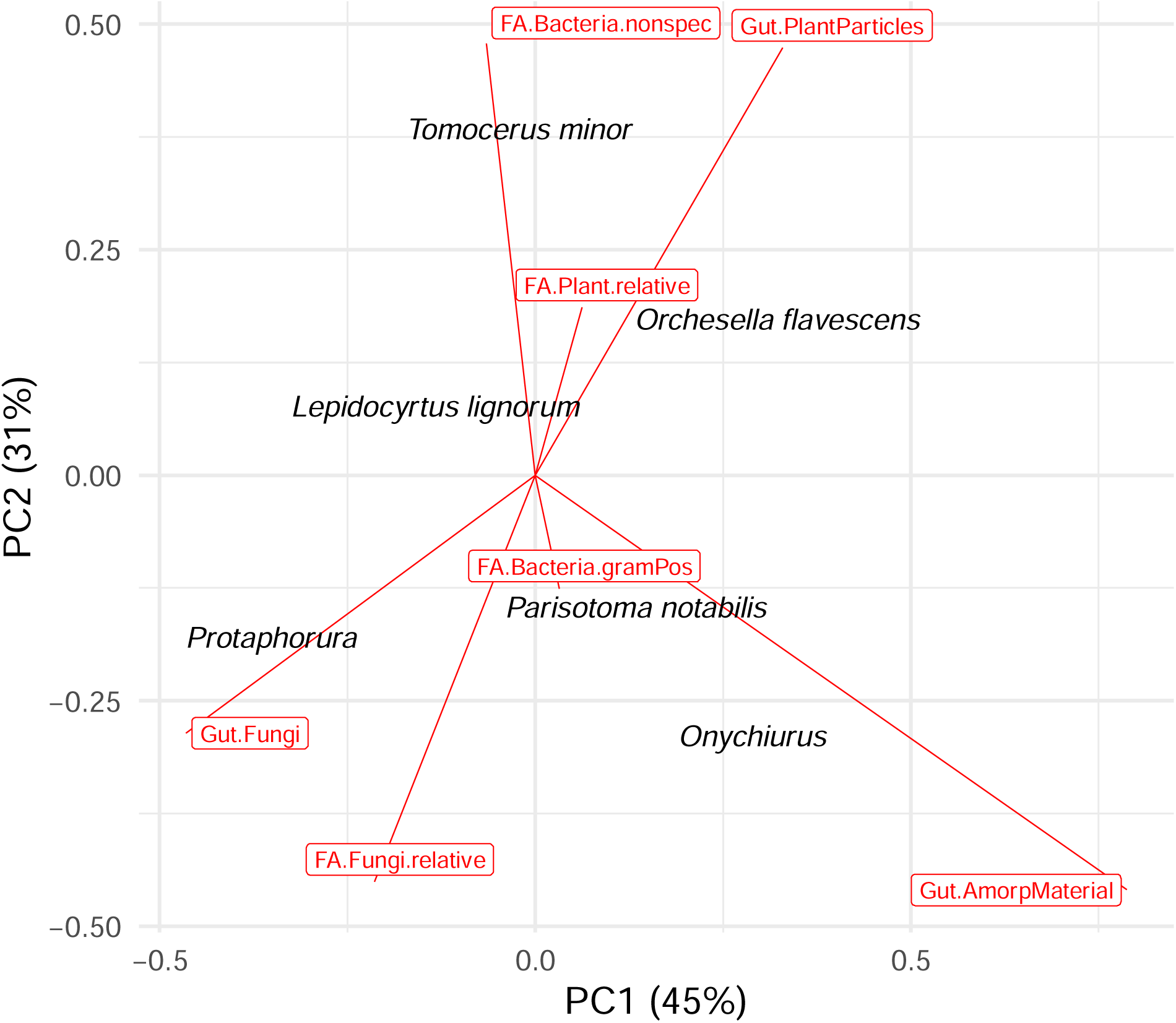
Principal component analysis based on seven selected trophic niche parameters of six common Collembola species. Species are shown with grey dots; trophic parameters are shown with red vectors.

### Taxonomic and life form effects on trophic niche parameters of Collembola

Taxonomic affiliation best explained stable isotope Δ^13^C and Δ^15^N, and also explained part of the variation in gram-positive and non-specific bacterial FA parameters, proportion of plant and fungi in the gut, and trehalase activity (all R^2^ ≥ 0.24, p < 0.05 except for trehalase **Fig. 3**). Cellulase activity and proportion of amorphous material in gut were poorly related to the taxonomic affiliation (R^2^ = 0.03).

**Figure 3.**
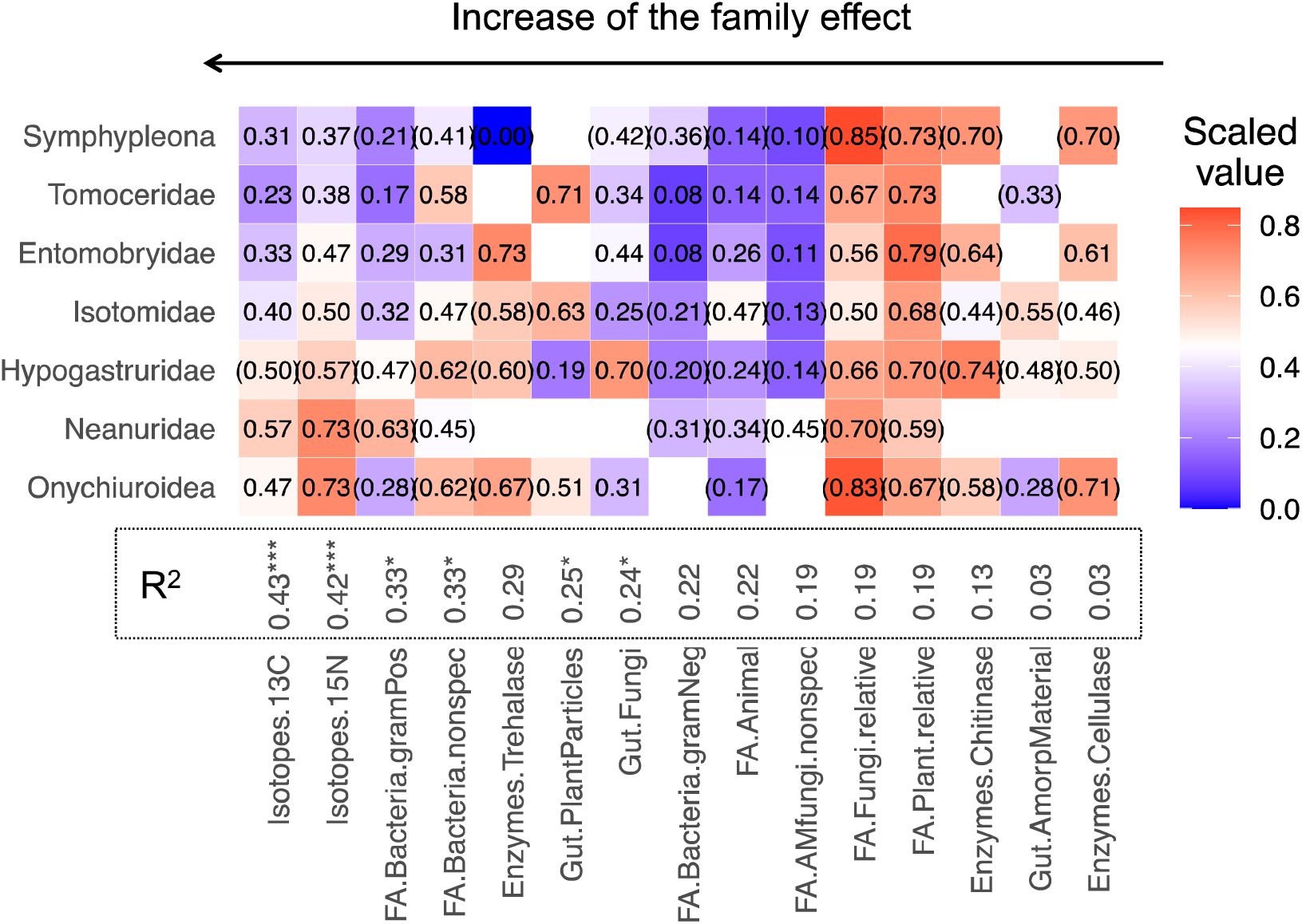
Median values of trophic niche parameters in Collembola taxonomic groups. All parameters were scaled between 0 (minimum, intense blue) and 1 (maximum, intense red). Parameter values based on < 3 species were excluded; values based on 3–5 species are given in brackets; in other cases, n = 6–20. Numbers next to parameters indicate R^2^ explained by taxonomic affiliation; ***p < 0.001, **p < 0.01, *p < 0.05. Taxonomic groups are sorted according to the orders; parameters are sorted according to the R^2^ values.

Analysed together, the seven Collembola taxonomic groups differed in their trophic niches (**Fig. 4**). Symphypleona had high average proportion of FAs synthesised by gram-negative bacteria (PC4) and fungi (PC2), high chitinase and cellulase (PC1), but very low trehalase activity (PC2). Tomoceridae were characterised by high average proportions of plant particles in the gut (PC1) and low Δ^13^C and Δ^15^N values (PC1, PC2). Entomobryidae had the highest trehalase activity (PC2) but low values of bacteria-related parameters (PC1). By contrast, Isotomidae had lower values of fungi-related parameters and enzyme activities (PC2, PC3, PC4). A further separation of Isotomidae into epedaphic and (hemi)edaphic life forms indicated that epedaphic Isotomidae had lower Δ^13^C values (0.35) than edaphic ones (0.49), a lower proportion of plants (epedaphic: 0.55; edaphic: 0.72) but a higher proportion of fungi in the gut (epedaphic: 0.42; edaphic: 0.20). Hypogastruridae had a remarkably high proportion of fungi in the gut and high chitinase activity, but a low proportion of plant particles in the gut (PC3). Neanuridae had the highest Δ^13^C and Δ^15^N values and proportions of FAs synthesized by gram-positive bacteria and arbuscular-mycorrhizal fungi (PC1). Onychiuroidea had high Δ^15^N values, FAs synthesized by non-specific bacteria and fungi, and high cellulase activity (PC4).

**Figure 4.**
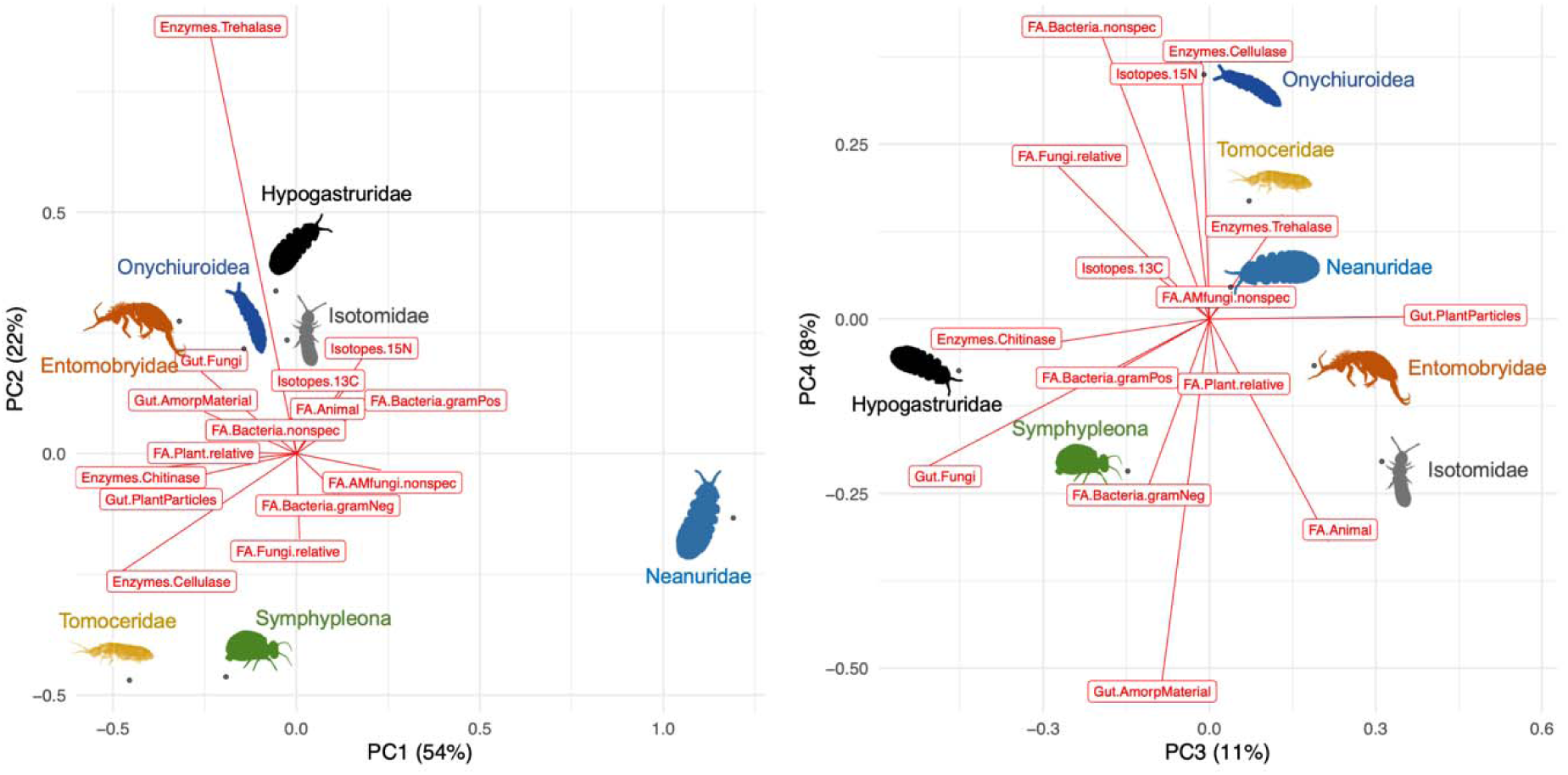
Principal component analysis based on median values of the fifteen trophic niche parameters of Collembola taxonomic groups. Collembola groups are shown with dots and silhouettes; trophic parameters are shown with red vectors. Most of the silhouettes were taken from http://phylopic.org, credit goes to Birgit Lang and Kamil S. Jaron.

Life form best explained chitinase activity, but the effect was not significant due to a low number of replicates. Life form also well explained Δ^15^N values and proportion of FA synthesised by gram-negative bacteria (both R^2^ ≥ 0.25; **Fig. 5**). Most of the other parameters were moderately or not at all related to life form.

**Figure 5.**
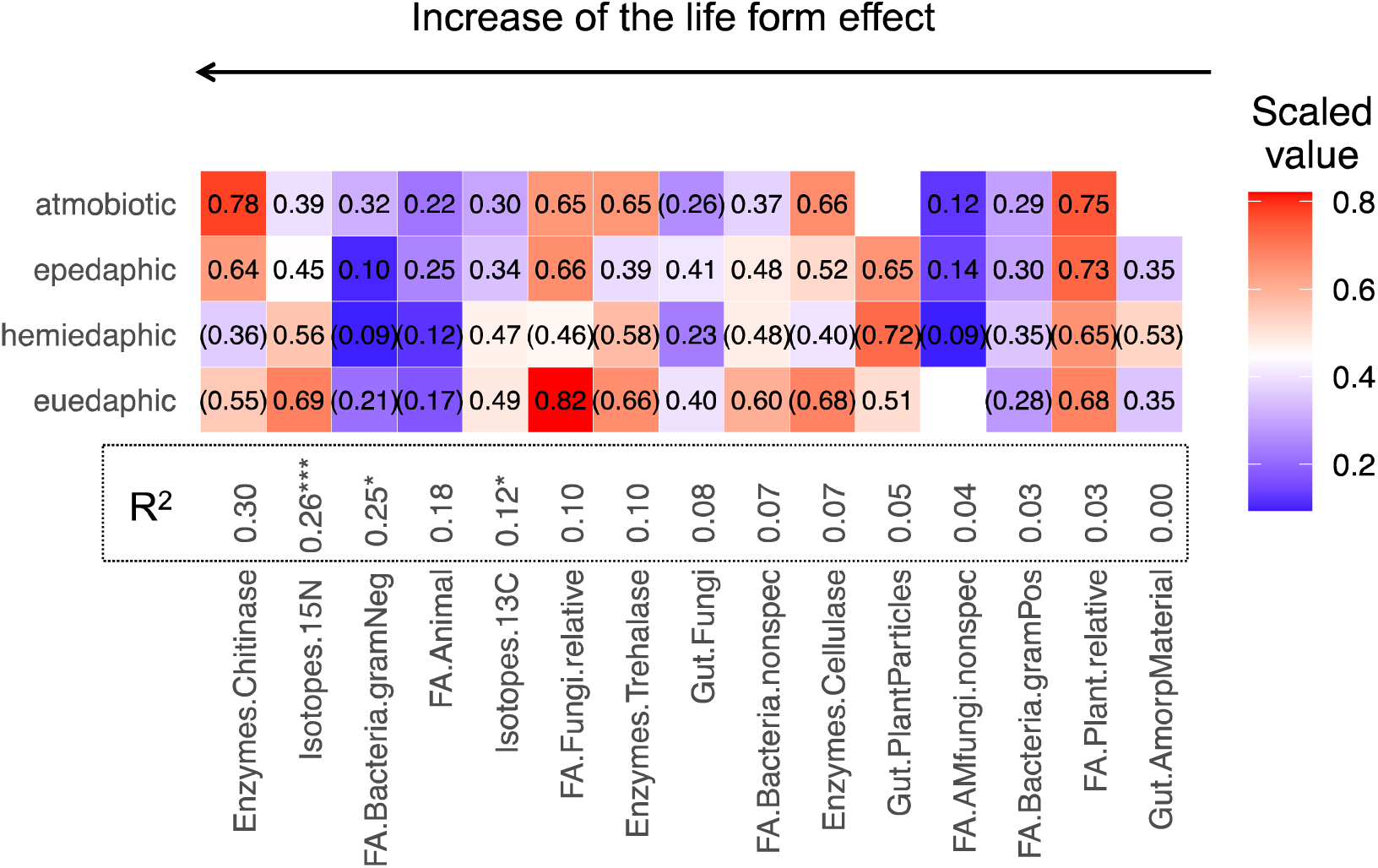
Median values of trophic niche parameters in Collembola life forms. All parameters were scaled between 0 (minimum) and 1 (maximum). Parameter values based on < 3 species were removed, values based on 3–5 species are given in brackets; in other cases, n = 6–36. Numbers next to parameters show R^2^ explained by life form identity; ***p < 0.001, **p < 0.01, *p < 0.05. Life forms are sorted according to the species vertical stratification along the soil profile; parameters are sorted according to the R^2^ values.

Atmobiotic species had high average activity of all digestive enzymes and a high proportion of FAs synthesized by gram-negative bacteria and plants. Epedaphic species had in general intermediate values for all parameters, but high proportions of fungi in the gut and high proportions of animal-synthesized FAs. Hemiedaphic species had high Δ^13^C values and proportion of FAs synthesized by gram-positive bacteria, but low activity of chitinase and cellulase. Euedaphic species had the highest Δ^15^N values, high trehalase activity, high proportions of FAs synthesised by fungi and non-specific bacteria biomarker FA.

## Discussion

Different dietary methods track different processes of animal feeding at different resolution and time scales, providing information on the multidimensional nature of trophic niches. However, studies quantitatively comparing different methods are scarce, and usually involve one or two methods only (Nielsen et al., 2018). Here, we compared fifteen trophic niche parameters derived from four methods using Collembola – a model group of the cryptic and diverse soil animals. First, we showed that trophic niche parameters were weakly correlated, reflecting complementarity, rather than redundancy, of different methods. Second, we outlined the trophic niche parameters that vary with taxonomic clusters and microenvironments of species. Finally, we presented the multidimensional trophic niche of Collembola, and provided the most detailed trophic information to date on this animal group.

### The additional value of method combination

We found weak correlations among trophic niche parameters derived from different methods in 70% of the performed tests. As already shown, outcomes provided by different dietary methods only in part overlap (Nielsen et al., 2018). Pessimistically, this reflects biases embedded in each method – optimistically, it means that different methods inform on more trophic niche dimensions (Hambäck, Weingartner, Dalén, Wirta, & Roslin, 2016). Our study demonstrated various types of interactions among the methods and parameters, including confirmation, controversy, complementarity and clarification. For instance, proportion of plant and fungal particles in gut correlated with retention of plant and fungal FAs, respectively. Visual gut content analysis and FA analysis confirm each other in detection of herbivory and fungivory in Collembola; applying one of the methods may sufficiently trace these feeding strategies. These two methods explore the trophic niches at different stages in dietary processes. Since visual gut content analysis detects ingested food particles and the FA method traces assimilated compounds, it also implies that in these two food categories, Collembola digest mostly the same material they ingest. Further, Δ^15^N and Δ^13^C values were negatively correlated with several plant-related parameters provided by the other methods, confirming (1) that high Δ^15^N values reflect high trophic level as evident from fewer plant particles in the gut, and (2) that high Δ^13^C values imply feeding on microbially-decomposed organic matter but not freshly-fixed plant carbon, thus having a lower need for cellulase (A. M. Potapov, Tiunov, & Scheu, 2019; A. M. Potapov, Tiunov, Scheu, Larsen, & Pollierer, 2019).

Controversy was found in fungi- or plant-related parameters between the digestive enzyme method and the FA method. Although we would expect positive correlations between chitinase activity (allowing to degrade chitin in fungal cell walls) and fungal FA proportions, or between cellulase activity (allowing to degrade cellulose in plant and algae cell walls) and plant FA proportions, neither was the case. To verify correlations of the fungal or plant parameters derived from the digestive enzyme method with those from visual gut content analysis, however, more data are needed.

Complementarity of the methods was shown in gut fungi and plant- and fungi-synthesized FAs that were positively correlated with trehalase activity. Since trehalose is the storage component of fungal, lichen and plant cells, these correlations suggest that Collembola feeding on plant material and fungi get energy and nutrients from storage, rather than structural, polysaccharides. In relation to the foraging behaviour, Collembola are more like browsers rather than grazers (Siepel & de Ruiter-Dijkman, 1993). Another example for complementarity of the methods was negative correlations of activity of trehalase and chitinase with the FAs synthesised by non-specific and gram-positive bacteria. Apparently, certain Collembola species have a specific feeding strategy where they rely more on bacterial feeding and thus invest less in the fungi-related digestive enzymes. Detritivores may largely rely on amino acids synthesised by free-living or gut symbiotic bacteria if the food quality is low (Larsen et al., 2016; A. M. Potapov, Tiunov, Scheu, et al., 2019). When repeatedly consuming decomposing litter and soil, soil animals may collaborate with microorganisms that produce digestive enzymes to release nutrients from recalcitrant organic compounds – a strategy termed “external rumen” (Swift, Heal, & Anderson, 1979). This is further supported by positive correlations of Δ^15^N values and FAs synthesised by non-specific and gram-positive bacteria, pointing to the utilization of bacterial symbionts when Collembola feed on soil organic matter. Higher Δ^15^N values in soil-dwelling Collembola species likely are due to trophic level inflation by repeated ingestion of soil organic matter (A. M. Potapov, Semenina, et al., 2016; Steffan et al., 2017). Gram-positive bacteria such as Actinobacteria and Firmicutes associate with small soil fractions and inhabit small pores (Hemkemeyer, Dohrmann, Christensen, & Tebbe, 2018; Mummey, Holben, Six, & Stahl, 2006). These bacteria are thus likely accessible for the soil feeders. Indeed, Actinobacteria dominate in the gut of *Folsomia candida*, a euedaphic Collembola species specifically adapted to the soil layer (Zhu et al., 2018).

We avoid to further discuss correlations since many of them were based only on few data points. In summary, our results showed that FA and visual gut content analyses both indicate herbivory and fungivory but reveal different stages of the dietary processes in Collembola. Digestive enzyme analysis has a high additional value when combined with others. It gives further insights in foraging behaviour and animal-microbial interactions: Collembola behave more like browsers, rather than grazers, by feeding on storage polysaccharides from plant material and fungi. Furthermore, stable isotope analysis estimates trophic level and plant versus microbial feeding of soil animals and the complementarity of these parameters and the other trophic parameters indicates that bacterial feeding in Collembola may be more common than usually assumed.

### Taxonomic and life form effects on trophic niche parameters

We further explored how the trophic niche parameters vary across different Collembola taxa and life forms. Taxonomic affiliation explained variation in six out of the fifteen trophic niche parameters, suggesting that some but not all trophic niche dimensions in Collembola are phylogenetically structured (A. M. Potapov, Semenina, et al., 2016). The taxonomic groups explained most variation in stable isotope and in bacteria-related FA parameters. These parameters are related to biochemical processes in assimilation and thus may in part be constrained by physiology of phylogenetically-related species (T.-W. Chen et al., 2017). By contrast, cellulase activity was not related to taxonomic groups, suggesting that the ability of cellulose degradation might be evolutionary labile in Collembola.

Life form, as a proxy for the microenvironment species live in, explained variation only in three parameters with relatively low R^2^ values. Most trophic niche parameters were poorly related to life form, suggesting that various feeding strategies may be used by species living in the same microenvironments. However, we found higher values of stable isotopes Δ^15^N and Δ^13^C in (hemi)edaphic species, suggesting that dwelling mainly in soil microhabitats results in feeding strategies that rely mainly on microbially-decomposed organic matter and less on plant materials. By contrast, higher chitinase activity in atmobiotic and epedaphic Collembola suggests that they need to digest fungal cell walls. Interestingly, atmobiotic species, which mainly inhabit macrophytes (e.g. on grasses, bushes, trunks and branches of trees), had higher proportions of gram-negative bacteria FAs, as compared to epedaphic species living in the upper layer of litter. Bacterial communities in these two microhabitats clearly differ from each other. Collembola consume gram-negative bacteria such as cyanobacteria (Buse, Ruess, & Filser, 2013; Hao, Chen, Wu, Chang, & Wu, 2020). The atmobiotic Collembola might feed on corticolous cyanobacteria and lichens associated with tree bark (A. Singh, Tyagi, & Kumar, 2017). Furthermore, even within the same Isotomidae family, differences in trophic niche parameters between epedaphic and (hemi)edaphic species point to environmental determinants of trophic traits in soil animals (Ponge, 2000; A. M. Potapov, Tiunov, & Scheu, 2019).

### Multidimensional trophic niches of Collembola

As practical demonstration of the additional value of method combination, below we attempt to describe the multidimensional trophic niches for high-rank Collembola taxa (**Fig. 3, 4**). The order Symphypleona (here representing a conglomerate of several surface-dwelling families) on average had high proportions of fungal FAs, high chitinase activity in fungal cell wall degradation, and a high proportion of FAs synthesised by gram-negative bacteria. Lichens are comprised of fungi and microalgae or cyanobacteria. Cyanobacteria are gram-negative bacteria able to synthesise hydroxylated FAs (Dembitsky, Shkrob, & Go, 2001; Gugger, 2002). Together with low Δ^15^N values typically found in algivores (A. M. Potapov, Korotkevich, & Tiunov, 2018), our results suggest that lichen grazing, or a combination of fungivory and herbivory may be widespread among Symphypleona.

Tomoceridae had a relatively well-defined trophic niche; the combination of parameters points to consumption of plant material with the help of bacteria. They had the highest average proportion of plant particles in the gut among all other Collembola and a relatively high proportion of plant-synthesised FAs. In addition, they had a high proportion of non-specific bacteria FAs, suggesting that they may graze on freshly fallen litter and assimilate it with the help of bacterial symbionts. Clarification of their trophic niche could be advanced with enzyme analysis, but only one record of *Tomocerus minor* was present in our database – this species had the highest cellulase activity among all records in the database, confirming plant litter grazing.

Entomobryidae include many species that are morphologically resembling Tomoceridae; however, these two families differed in a number of trophic niche parameters. Entomobryidae had more fungi in the gut, remarkably high trehalase activity, but lower proportion of non-specific bacteria FAs. Overall, these differences suggest that Entomobryidae are likely browsers, rather than grazers. *Lepidocyrtus lignorum* is a good illustration – this species preferred fungi (**Fig. 2**), having high trehalase, but low cellulase and limited chitinase activity (**Table S2**).

Isotomidae had lower average values of fungi-related parameters and lower enzymatic activity than the previous two families. Isotomidae also had high Δ^13^C values, pointing to consumption of organic material in advanced stages of decomposition (A. M. Potapov, Tiunov, & Scheu, 2019), potentially including invertebrate faeces. All these features were the most expressed in hemiedaphic and euedaphic species that inhabit decomposing litter and soil (e.g. *Parisotoma notabilis*; **Fig. 2**). It is likely, that edaphic Isotomidae species rely more on bacteria via using the “external rumen” feeding strategy. They are usually small and their gut often contains humus (i.e. soil and faecal material) (Ponge, 2000; Poole, 1959). Interestingly, hemiedaphic and euedaphic Isotomidae such as species of the genera *Isotomiella, Folsomia, Parisotoma, Folsomides*, dominate in Collembola communities in many ecosystems (M. Potapov, 2001). Bacterial feeding may be more common in Collembola than it is assumed in traditional soil food web models (de Vries et al., 2013; Hunt et al., 1987).

Hypogastruridae had the highest proportion of fungi and the lowest proportion of plants in the gut across all other Collembola and low activity of cellulase, suggesting that they feed selectively on microorganisms and not on plant material. Species of this family were reported to live and feed on fungi (Sawahata, Soma, & Ohmasa, 2001), and to have relatively high proportions of fungi-synthesised FAs (Ferlian et al., 2015). Taken together, Hypogastruridae are fungivores, or, more broadly, microbivores.

Neanuridae have no molar plate and are thus unable to chew the food. This family was long recognised to have a distinct trophic niche from other families (Berg et al., 2004; Chahartaghi et al., 2005; S. B. Singh, 1969). However, their exact food objects remain enigmatic. Early studies hypothesised that Neanuridae feed by sucking up the content of fungal hyphae (Poole, 1959; S. B. Singh, 1969). Stable isotope analysis discovered that Neanuridae have an outstandingly high trophic level, suggesting that they may feed on other animals, such as nematodes (Chahartaghi et al., 2005). More recently, Neanuridae were shown to successfully breed on slime moulds (protists with mycelial stage) (Hoskins, Janion-Scheepers, Chown, & Duffy, 2015). Slime moulds are abundant in various ecosystems (Swanson, Vadell, & Cavender, 1999) and tend to live in the rotten wood where Neanuridae are also found. Slime moulds, therefore, potentially serve as food for some species of this family in natural environments. In our study we showed that Neanuridae have high average values of bacteria- and fungi-related parameters, high Δ^13^C and Δ^15^N values, but very low plant supplementation, supporting the abovementioned hypotheses. Interestingly, Neanuridae also had a high proportion of FAs synthesized by arbuscular mycorrhizal fungi; however, the same biomarkers can also be synthesized by bacteria. Neanuridae, as high-level consumers among Collembola, feed on various groups of microfauna and microorganisms and receive energy from both bacterial and fungal origins.

Onychiuroidea are soil-adapted Collembola without eyes, pigment and furca. They are associated with plant roots, presumably by feeding on root tips or mycorrhizae (Endlweber, Ruess, & Scheu, 2009; Fujii et al., 2016; A. M. Potapov, Goncharov, Tsurikov, Tully, & Tiunov, 2016). They had intermediate proportions of plant and fungal particles in their gut in comparison to other Collembola. However, they had high proportions of FAs synthesised by fungi and non-specific bacteria and a high cellulase activity. Potentially, Onychiuroidea may feed on both roots and soil. However, the small-sized Tullbergiidae, which were analysed with Onychiuridae in our study, may rely less on the root-derived resources (Li et al., 2020). Food resources of this superfamily call for more studies.

Using comparisons across multiple methods, we provide a general overview of Collembola trophic niches. Our study is the first cross-method compilation of available data on trophic traits in Collembola that clearly shows advantages of the multidimensional trophic niche approach. When combined, different methods are not redundant but rather have a high additional value by compensating drawbacks of each other. Simultaneous application of several methods to the same population across different groups and ecosystems may improve our understanding of functioning of the food webs and help to explain species coexistence in cryptic environments, such as soil.

## Acknowledgements

This study was funded by the Russian Science Foundation (project 19-74-00154) to AMP. Joint discussions among TWC, VS, MMP and AMP were supported by CAS-DAAD mobility project (DAAD-19-10) co-funded by Czech Academy of Sciences (CAS) and German Academic Exchange Service (DAAD, Deutscher Akademischer Austauschdienst). TWC was supported by the MSM project funded by CAS for research and mobility support of starting researchers (MSM200962001). MMP was supported by the German Research Foundation (DFG, MA 7145/1-1).

## Authors’ contributions

AMP and TWC developed the idea and study design. SS compiled data on the gut content analysis, VS compiled data on the digestive enzyme analysis, MMP compiled data on the fatty acid analysis, AMP compiled data on the stable isotope analysis. AMP and TWC did the analysis and drafted the manuscript. All authors critically revised the analysis and text.

## Data accessibility

Raw data supporting the results of the study are provided in the Supplementary materials (Table S1-4) and we intend to archive it to the Dryad digital repository, should the manuscript be accepted for publication.

## Supplementary materials

- Table S1 – dataset, gut content data (provided separately)
- Table S2 – dataset, enzyme data (provided separately)
- Table S3 – dataset, FA data (provided separately)
- Table S4 – dataset, stable isotope data (provided separately)
- Table S5 – overlap between different trophic-niche parameters
- Figure S1 – pairwise correlation between trophic-niche parameters, full version
- Reference list for the Supplementary materials

## Notes

### Competing Interest Statement

The authors have declared no competing interest.

