## Supplementary materials for "Multidimensional trophic niche revealed by complementary approaches: gut content, digestive enzymes, fatty acids and stable isotopes in soil fauna"

- Table S1 – dataset, gut content data (provided separately)
- Table S2 – dataset, enzyme data (provided separately)
- Table S3 – dataset, FA data (provided separately)
- Table S4 – dataset, stable isotope data (provided separately)
- Table S5 – overlap between different trophic-niche parameters
- Figure S1 – pairwise correlation between trophic-niche parameters, full version
- Reference list for the Supplementary materials

**Table S5**. Overlap between different trophic-niche parameters. The numbers of species that has records on each of two parameters for each pairwise comparison are given.

|  | FA.Bacteria.gramPos | FA.Bacteria.gramNeg | FA.Bacteria.nonspec | FA.AMfungi.nonspec | FA.Fungi.relative | FA.Plant.relative | FA.Animal | Gut.PlantParticles | Gut.Fungi | Gut.AmorpMaterial | Enzymes.Trehalase | Enzymes.Cellulase | Enzymes.Chitinase | Isotopes.13C | Isotopes.15N |
| --- | --- | --- | --- | --- | --- | --- | --- | --- | --- | --- | --- | --- | --- | --- | --- |
| FA.Bacteria.gramPos | 41 |  |  |  |  |  |  |  |  |  |  |  |  |  |  |
| FA.Bacteria.gramNeg | 34 | 34 |  |  |  |  |  |  |  |  |  |  |  |  |  |
| FA.Bacteria.nonspec | 41 | 34 | 47 |  |  |  |  |  |  |  |  |  |  |  |  |
| FA.AMfungi.nonspec | 33 | 33 | 33 | 33 |  |  |  |  |  |  |  |  |  |  |  |
| FA.Fungi.relative | 41 | 34 | 47 | 33 | 47 |  |  |  |  |  |  |  |  |  |  |
| FA.Plant.relative | 41 | 34 | 47 | 33 | 47 | 47 |  |  |  |  |  |  |  |  |  |
| FA.Animal | 36 | 29 | 40 | 29 | 40 | 40 | 40 |  |  |  |  |  |  |  |  |
| Gut.PlantParticles | 6 | 2 | 6 | 2 | 6 | 6 | 5 | 40 |  |  |  |  |  |  |  |
| Gut.Fungi | 11 | 7 | 11 | 7 | 11 | 11 | 9 | 40 | 54 |  |  |  |  |  |  |
| Gut.AmorpMaterial | 7 | 3 | 7 | 3 | 7 | 7 | 6 | 32 | 34 | 34 |  |  |  |  |  |
| Enzymes.Trehalase | 10 | 9 | 10 | 9 | 10 | 10 | 9 | 3 | 7 | 3 | 24 |  |  |  |  |
| Enzymes.Cellulase | 10 | 9 | 10 | 9 | 10 | 10 | 9 | 3 | 5 | 3 | 22 | 22 |  |  |  |
| Enzymes.Chitinase | 10 | 9 | 11 | 9 | 11 | 11 | 10 | 3 | 7 | 3 | 22 | 20 | 23 |  |  |
| Isotopes.13C | 22 | 17 | 24 | 17 | 24 | 24 | 22 | 14 | 19 | 11 | 14 | 13 | 12 | 90 |  |
| Isotopes.15N | 22 | 17 | 24 | 17 | 24 | 24 | 22 | 14 | 19 | 11 | 14 | 13 | 12 | 88 | 88 |

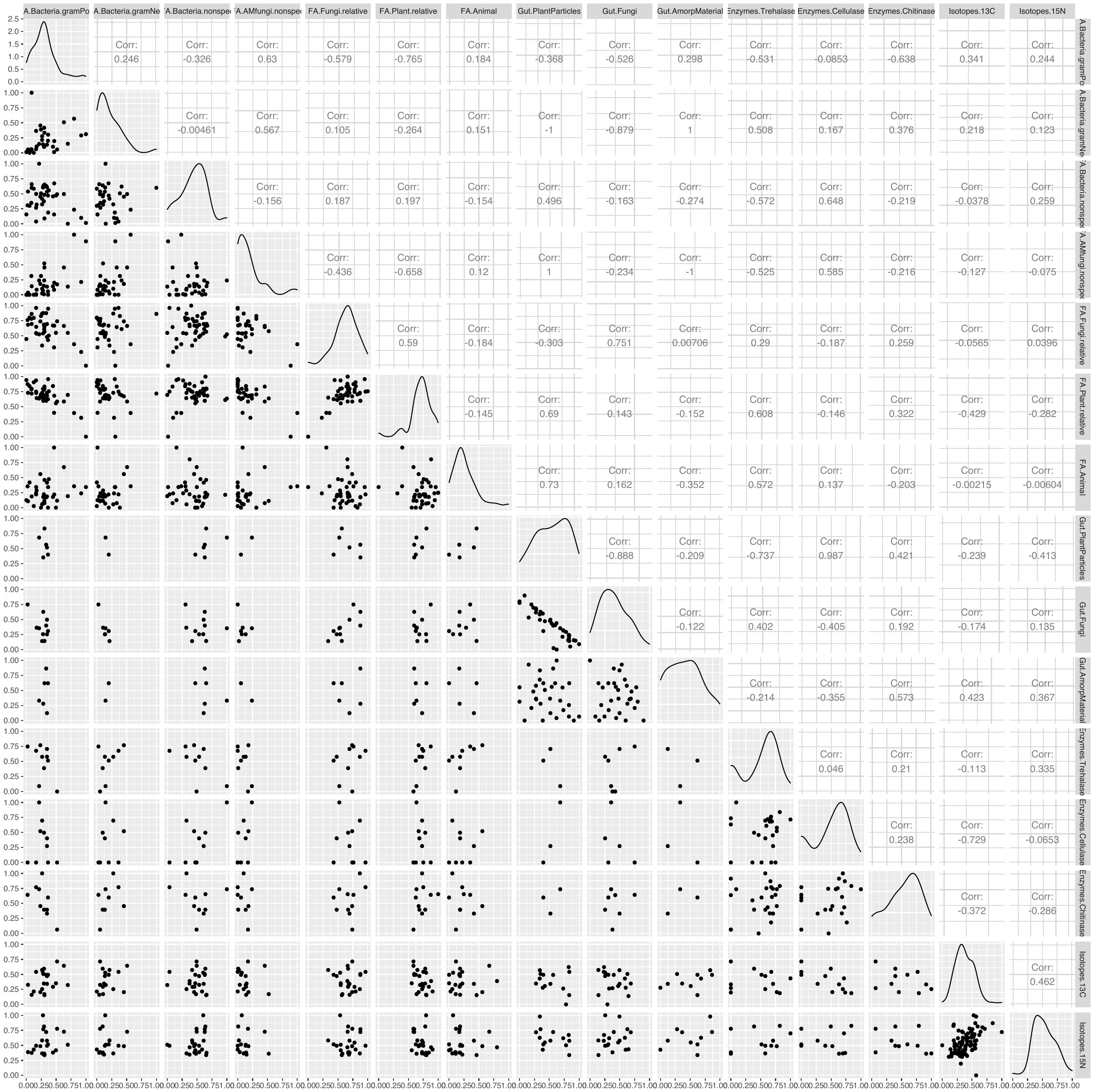

**Figure S1.** Correlation among fifteen trophic niche parameters in Collembola produced with *ggpairs* in *GGally* package. Number of points varies from 0 (“NA”) to 88, depending on the method overlap. Spearman correlation coefficients (“Corr:”) are shown in the upper triangle. Diagonal shows density of the data distribution for corresponding parameters. Figure provides more detailed view of the analysis presented on the Fig. 1 in the main text.
